# HEMIBIOTROPHIC INFECTION OF SUNFLOWER ROOT CELLS BY THE PARASITE *OROBANCHE CUMANA* WALLR

**DOI:** 10.1101/2024.10.08.617216

**Authors:** S.H. Khablak, V.M. Spychak, Y A. Abdullaieva, Ya.B. Blume

## Abstract

The process of infection of different sunflower hybrids by Orobanche cumana was studied according to the way the pathogen interacts with the components of root cells. The study of the behavior of sunflower root cells during the penetration of Gaustoria showed not biotrophic, but hemibiotrophic process of infection of hybrids with the pathogen. In the hemibiotrophic mixed type of sunflower infection, broomrape initially develops as a biotrophic pathogen and, with the help of effectors, suppresses the course of immune reactions and forms multicellular haustoria, which at a certain stage of growth break the membrane bags of cells, leading to cytoplasmic leakage and causing the death of root cells as a necrotrophic pathogen. The discovery of the hemibiotrophic process of sunflower broomrape infection opens a new promising direction in reducing the infection of hybrids with the parasite through the induction of systemic acquired resistance (SAR) by drugs that cause the formation of reactive oxygen species and trigger plant defense responses through programmed cell death at the sites of infection and cause pathogen necrosis. The process of infection of cells and the emergence of reverse immune responses in plants when infected with pathogens has similar features and a similar course of protective responses. In the interaction between broomrape and sunflower, resistance mechanisms may act to stop the pathogen in the root cortex, in the entodermis, or after reaching the central cylinder. The cell wall is the first obstacle that pathogens have to overcome. The cell membranes of growing cells have a “primary” structure. Cell walls are composed of cellulose and matrix substances (hemicelluloses, pectins, and proteins). In cells that have already formed, the cell walls are reinforced with lignin, suberin, and kalose. Sunflower’s defense reactions against broomrape consist of strengthening the cell wall through the deposition of lignin, suberin, accumulation of callose, protective proteins and cross-linking of proteins that prevent the parasite from penetrating and communicating with the host vascular system. Lignin, suberin and callose polymers strengthen the cell wall structure by increasing its stiffness at the site of infection to limit the penetration of the parasite.

## Introduction

The broomrape (Orobanche cumana Wallr.) is an important parasite in sunflower crops in Europe, some Asian countries and Australia, and Ukraine (Martín-Sanz et al., 2016). It is estimated that 16 million hectares of sunflower in Europe and Asia, especially in Central and Eastern Europe, Spain, Turkey, Israel, Iran, Kazakhstan, Ukraine and China are infested with broomrape. Annual yield losses worldwide as a result of broomrape infestation of sunflower areas are estimated at approximately 1.17-2.33 billion euros (Cvejić et al., 2020).

In recent years, Ukraine has been experiencing broomrape infestation of sunflower hybrids that are resistant to races E, F and G. From the northern Steppe of Ukraine, the broomrape is actively moving to the central, northern and western regions of the country. This is due to the emergence of new centers and races of the parasite in these areas. Today, the problem of broomrape is of global importance (Khablak, Spychak, 2023).

According to current research statistics, the best way to reduce the damage caused by broomrape is to develop resistant sunflower hybrids. To do this, it is necessary to find out the cellular and molecular mechanisms of crop resistance to the parasite, which will be useful in developing new effective approaches to control parasitic plants (Sisou et al., 2021).

Plant pathogens are classified according to the way they feed. Necrotrophic pathogens actively kill host tissue when they colonize and thrive on the contents of dead or dying cells. This way of life is different from that of biotrophic pathogens, which obtain nutrients from living cells and therefore must keep the host viable. This difference underlies the diverse pathogenesis strategies and immune responses of plants to biotrophic and necrotrophic infections. The third group, hemibiotrophs, demonstrate both forms of nutrient acquisition, switching from an early biotrophic phase to necrotrophy in the later stages of the disease. The duration of the biotrophic or necrotrophic phase varies considerably among hemibiotrophic pathogens (Laluk, Mengiste, 2010).

Despite the taxonomic differences between parasitic species, all parasitic plants have haustoria, including the parasite Orobanche cumana. A haustoria is a unique, specialized organ that originates from lateral roots and allows pathogenic fungi and parasites to parasitize other plants. The word haustorium is derived from the Latin “haustor or haurire”, which means “water box”. At different stages of development, the haustorium functions in host attachment, host invasion, evasion of host immunity, and nutrient transfer (Yoshida et al., 2016).

The fungal haustoria is a unicellular hyphae, while the haustoria of parasitic plants is a multicellular organ. The fungal haustorium is an intracellular or intercellular structure that infects plants, depending on the pathogen, intracellularly and is surrounded by the host extrahaustorial membrane or intercellularly, while the parasitic plant haustorium is an intercellular structure that penetrates between host cells (Mendgen, Hahn, 2002).

Intercellular penetration of Gaustoria was shown during the infection of cowpea (Vigna unguiculata) with the parasite Striga gesnerioides (family Orobanchaceae) (Reiss and Bailey, 1998). In the stem parasitic species of Cuscuta (Convolvulaceae), intracellular and intercellular penetrations were found during infection of Pelargonium zonale (Press et al., 1990). There is evidence of intracellular penetration of intrusive cells of the parasitic plant Agalinis aphylla from the family Orobanchaceae into host cortical cells through a small hole in the cell wall (Musselman and Dickison, 1975).

It was believed that the intrusive cells at the tip of the haustoria of the broomrape parasite penetrate the sunflower root tissue intercellularly. During the process of penetration of the haustoria, the cell walls of sunflower cortical cells are dissolved by enzymes of the intrusive cells. In addition to the dissolution of the cell walls of sunflower root cells, the mechanical pressure from the penetrating intrusive cells of the parasite pushes the host cells aside so that the cell shapes change and the space between them is completely occupied by the pathogen’s haustoria (Perez-de-Luque, 2013).

Recent studies have shown that the broomrape is a biotrophic parasite and its penetration into sunflower root tissue occurs intracellularly, not intercellularly. When sunflower is infected, intrusive cells of the parasite Gaustoria penetrate intracellularly into the root epidermal and cortical layers, cross the entodermis and pericycle and reach the vessels of the host xylem and phloem, where they establish connections with them to absorb nutrients and water. As a result of local enzymatic decomposition and mechanical pressure exerted by intrusive Gaustoria cells while moving to the vessels, crossing the tissues in succession, deformation occurs, degradation of the cell wall in the root cells and pressing the plasmalemma into the middle of the cells, which is not destroyed. As a result of this process, a membrane sac is formed, which contains the parasite’s haustoria (Auriac et al., 2024).

It is known that when plants are infected with biotrophic pathogens, unlike necrotrophic pathogens, the spread of Gaustoria is ahead of necrosis. Gaustoria releases into the plant not toxins that kill cells, but effectors that suppress the course of immune reactions. The pathogen secretes some enzymes from the tip of the haustoria, which is in contact with the plant cell wall, that locally destroy the cell wall, and through the haustoria opening reaches the plasmalemma, which it does not destroy but gently presses into the cell interior. A membrane sac is formed, in which the extended end of the hypha, the haustoria, is located. At the same time, biotrophic pathogens do not have plasmalemma ruptures. Due to the absence of cytoplasmic membrane ruptures, the cell contents do not spill into the intercellular space and do not become a substrate for the parasite’s nutrition, as is the case with necrotrophic fungi (Shao et al., 2021).

However, cellular processes of sunflower broomrape infection involved in the penetration of intrusive Gaustoria cells into the root tissues of hybrids and the way they interact with cell components remain poorly studied and described. Our observations have shown that at a certain stage of infection of sunflower root with broomrape, when Gaustoria penetrates into the cells and forms a membrane sac and further growth of invasive parasitic structures, membrane ruptures and cytoplasmic leakage are observed, indicating not biotrophic but hemobiotrophic process of infection of the host with the pathogen. This requires research into the way intrusive Gaustoria broomrape cells interact with sunflower root cell components to confirm this fact.

In view of this, the aim of the research was to study the process of infection of sunflower root cells by the parasite Orobanche cumana according to the way the pathogen interacts with cell components. This knowledge is needed for further research on the cellular mechanisms of sunflower resistance to the parasite and the development of effective control measures for this parasite.

### Research methodology

The object of research in the vegetation experiments was broomrape seeds. Samples of the parasite seeds were collected on some of the most infected sunflower fields in the Forest-Steppe and Polissya. To study the process of sunflower broomrape infestation, hybrids of Lidea breeding company were used: ES Nirvana, ES Romantic, ES Genesis, ES Bella, ES Andromeda, ES Janis, ES Niagara, ES Artik.

The determination of the phenological stages of Orobanche cumana, at which infection occurs or resistance of sunflower hybrids to the pathogen occurs, was carried out using the rhizotron method (transparent plexiglass boxes), which allows for several weeks to monitor broomrape on sunflower roots and observe early stages such as compatible/incompatible attachments, tubercle development and tubercle necrosis (Le Ru et al., 2021).

Sunflower seedlings infected with O. cumana were grown in rhizotrons consisting of transparent plexiglass boxes containing a layer of mineral wool and paper covered with nutrient solution. Unlike growing in the field, the use of rhizotrons for cultivation of sunflower infected with O. cumana allows to observe the early stages of interaction between the parasitic plant and its host from the induction of germination of pathogen seeds to the tubercle stage. Resistance in sunflower samples can be characterized at the stage before attachment to the host, at the stage of attachment before the formation of haustoria (compatible/incompatible) and at the stage of tubercles after the formation of haustoria (quantification of the number of tubercles and necrosis of tubercles). The appearance of tubercles was defined as the period after the formation of haustoria and the establishment of vascular connections. The number of tubercles on the sunflower roots allows to distinguish susceptible and resistant sunflower genotypes at an early stage in the post-gaustorial period.

Observations of the parasite penetration before attachment to the host and during the pre-gaustorial period were carried out from 4 to 10 days after infection. Lepidoptera rarely penetrates the host root before 6 days, while most gaustoria reach the internal tissues of the root (inner cortex to blood vessels) after 8 days. The first attachments and first tubercles were visible 8 days and 15-20 days after infection, respectively. The development of buds from tubercles could be observed after one month of cultivation. Variability in the number of tubercles was observed after three weeks of cultivation in the resotron.

Sunflower roots were examined in the post-gastrointestinal period on days 14, 21, 28, and 35 after infection under a stereoscopic microscope “MBS-10” to determine the total number and stage of development of attachments to sunflower roots and the number of necrotic attachments. The stages of determining the phenological stages of Orobanche cumana at which infection occurs or resistance of sunflower hybrids occurs were based on the following classification with minor changes (Martín-Sanz et al., 2016). The following developmental stages were used: T00: no germination of broomrape seeds, T0: no attachment; T1: attachment formed, but the actual tubercle is not yet visible; T2: tubercle less than 1 mm in diameter; T3: tubercle larger than 1 mm in diameter and without visible stem buds; T4: tubercle with already formed stem buds or early stages of stem growth. T00 - stage before attachment to the host, T0, T1 - pregaustorial stage, T2, T3, T4 - postgaustorial period.

Cytochemical methods combined with light microscopy were used to study the process of infection of different sunflower hybrids by Orobanche cumana depending on the way the pathogen interacts with root cell components. Pieces of sunflower root with attached O. cumana seedlings were cut from sunflower plants grown by the rhizotron method using a stereoscopic microscope “MBS-10”.

The samples were prepared as follows. Half of the samples were fixed in a mixture of ethanol: acetic acid (3:1 by volume) for 10 minutes, cleared in 5 g/ml chloral hydrate for 48 hours with stirring, and examined using a Bresser Biolux LCD 50x2000x digital microscope and a Levenhuk MED 45T microscope, which allows for “dry” or immersion observation, using dark or bright field, phase contrast microscopy, or Koehler illumination. Phase-contrast microscopy increases the contrast and clarity of translucent and transparent samples to a level that can only be achieved by staining in classical studies (Pérez-de-Luque et al., 2008).

The rest of the samples were fixed in FAA solution (10% formaldehyde, 5% acetic acid, and 50% ethanol) for 5 minutes, dehydrated in a series of ethanol (50, 80, 95, 100, 100%: 12 hours each), and embedded in paraffin. Then, thin sections of 7-10 μm thickness were made using a Reichert-Jung 2040 microtome, stained with 0.2% toluidine blue for 3 minutes and examined under a Bresser Biolux LCD 50x2000x digital microscope. This method can detect phenolic substances as well as lignin and suberin. It also works well as a general stain, but in combination with other microscopic techniques it can provide valuable information.

### Research results and discussion

The degree of sunflower hybrids damage by broomrape is presented in Table 1. The results obtained indicate that all sunflower hybrids were affected by the parasite. No sunflower hybrids with resistance to Orobanche cumana were found.

**Table 1:** Degree of sunflower hybrids damage by broomrape.

| Hybrid | Selection | Resistance to broomrape | Number of plants tested, pcs. | Affected plants, % | Degree of broomrape damage | Number of broomrape nodules per 1 affected plant (average) |
| --- | --- | --- | --- | --- | --- | --- |
| EU Nirvana | Lidea | A-G | 20 | 80 | weak | $3 \pm 0,2$ |
| EU Romantic | Lidea | A-G | 20 | 82 | weak | $2,5 \pm 0,3$ |
| EU Genesis | Lidea | A-G | 20 | 75 | weak | $2,0 \pm 0,4$ |
| EU Bella | Lidea | A-G | 20 | 78 | weak | $2,6 \pm 0,3$ |
| EU Andromeda | Lidea | A-G | 20 | 74 | weak | $2,8 \pm 0,3$ |
| EU Janis | Lidea | A-G | 20 | 81 | weak | $2,0 \pm 0,2$ |
| ES Niagara | Lidea | A-G | 20 | 79 | weak | $2,3 \pm 0,3$ |
| ES Artik | Lidea | A-G | 20 | 77 | weak | $2,4 \pm 0,2$ |
| HIP<sub>05</sub> |  |  |  |  |  | 0,6 |

Notes: broomrape infestation of 7 or more nodules per 1 affected plant (average value) (7-10 points) is severe; 4-6 nodules (4-6 points) is moderate; 1-3 nodules (1-3 points) is weak.

The process of infection of cells and the emergence of reverse immune responses in plants when infected with pathogens has similar features and a similar course of protective responses. In the interaction between broomrape and sunflower, resistance mechanisms may act to stop the pathogen in the root cortex, in the entodermis, or after reaching the central cylinder. The cell wall is the first obstacle that pathogens have to overcome. The cell membranes of growing cells have a “primary” structure. Cell walls are composed of cellulose and matrix substances (hemicelluloses, pectins, and proteins). In cells that have already formed, the cell walls are reinforced with lignin, suberin, and callose.

Sunflower’s defense reactions against broomrape consist of strengthening the cell wall through the deposition of lignin, suberin, accumulation of callose, protective proteins and cross-linking of proteins that prevent the parasite from penetrating and communicating with the host vascular system. Lignin, suberin and callose polymers strengthen the cell wall structure by increasing its stiffness at the site of infection to limit the penetration of enzymes secreted by the pathogen.

As a rule, the penetration of broomrape with pre-gaustoral resistance stops in the sunflower root cortex on day 7-10 and is associated with darkening of the parasite seedlings. With post-gaustorial resistance, the pathogen movement is inhibited in the endodermis or after reaching the central cylinder on day 15-20 and causes necrosis of tubercles, which prevents the establishment of effective vascular connections with the host due to the occurrence of cell wall thickening in phloem cells and xylem vessels due to the accumulation of lignin, suberin, kalose, formation of protective proteins PR, antimicrobial substances, secondary metabolites such as phenolic compounds, recognition of the pathogen by resistance proteins R (leucine-rich NB-LRR or NLR) in the cytoplasm of the cell, which leads to ETI immunity and cell death. The penetration of broomrape in post-gaustorial sunflower resistance stops due to the strengthening of the cell wall not in the root cortex, but deeper in the entoderm or after reaching the central cylinder due to lignification of the host’s entodermal and pericyclic cells, which prevents the parasite from entering the root vascular cylinder.

All the sunflower hybrids tested were not resistant to the parasite. The broomrape haustoria established effective vascular connections with the host and further developed into a thickening that appeared on the root of the host plant, which was covered with tubercles that gave it the appearance of an asterisk. Subsequently, a bud formed at the opposite end of the asterisk, which was covered with numerous scales that later turned into modified leaves. Later, the bud developed into a flower-bearing stem that brings the inflorescence to the soil surface. The stages of sunflower broomrape infestation are very precise and adjusted in time. The first attachments on the roots of the broomrape occurred on day 7-10, and the formation of tubercles was observed on day 15-20. The development of buds from the tubercles could be observed after one month of cultivation.

To elucidate the cellular processes of sunflower infection by the broomrape parasite, we studied the penetration of intrusive Gaustoria cells into the root tissue and damage to cell components using microscopy approaches (Fig. 1). The studies showed that intrusive Gaustoria cells make their way to the host vessels, crossing the sunflower root tissue sequentially. At the same time, broomrape haustoria penetrates into the living tissues of the sunflower root as a result of degradation of the host cell wall and formation of a membrane sac. However, at the further stage of infection with Orobanche cumana, when Gaustoria penetrates into the cells and forms a membrane sac and subsequent growth of invasive parasitic structures, membrane ruptures and cytoplasmic leakage are observed, indicating not biotrophic but hemibiotrophic process of infection with sunflower pathogen.

**Fig. 1.**
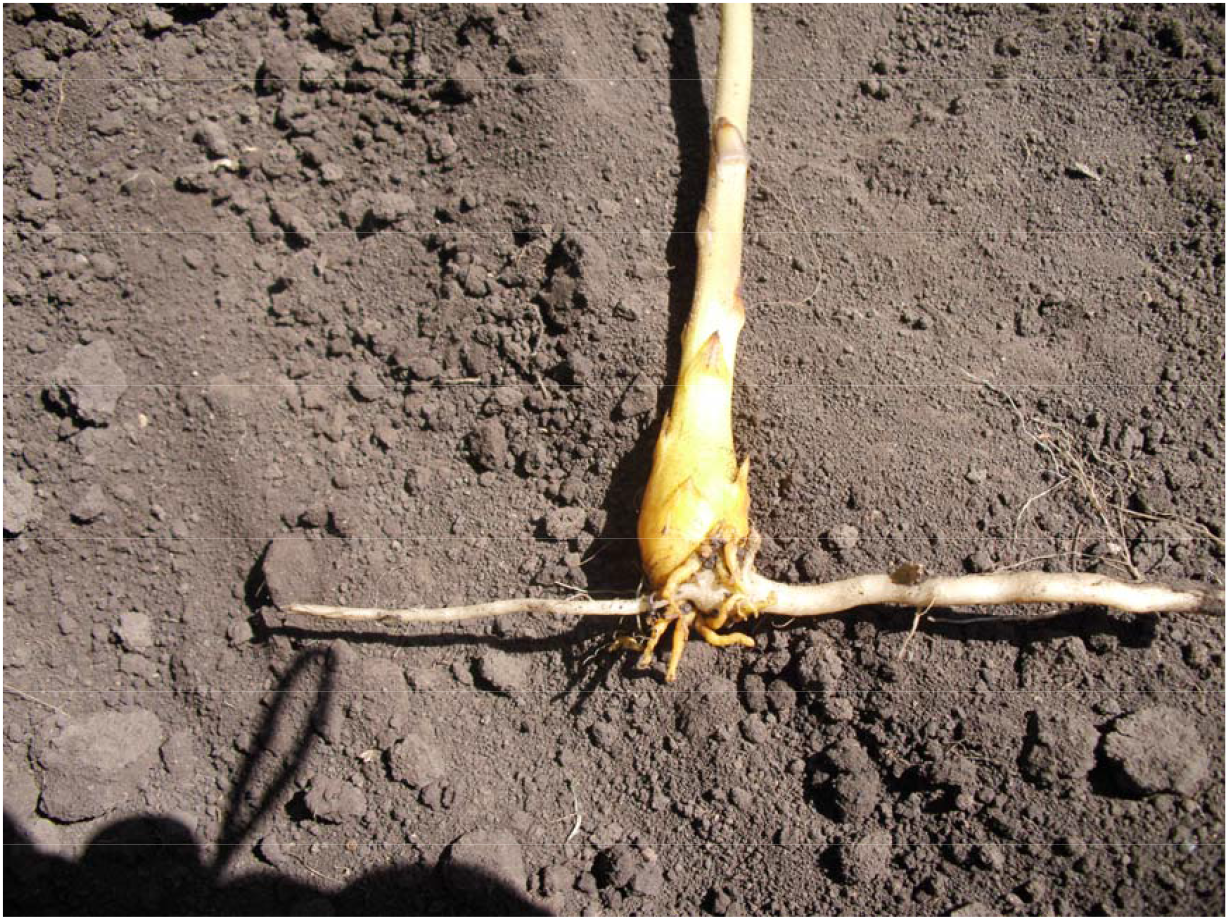
Gaustoria broomrape on sunflower roots.

The broomrape seed accepts its host thanks to germination stimulants present in sunflower root exudates. After germination, the root of the broomrape grows towards the host root and develops papillae that adhere to the sunflower root and secrete mucilaginous compounds. Over time, epidermal cells at the tip of the haustoria, a specific parasitic organ, differentiate into intrusive cells that penetrate the host root. This penetration combines physical pressure and degradation of the sunflower root cell walls due to pectolytic enzymes released by the parasitic plant. The intrusive cells make their way to the vessels of the host root, crossing the sunflower tissue sequentially. After contact with the host xylem vessels, the intrusive cells differentiate into vascular elements and vascular connections are established (xylem as well as phloem) to ensure the supply of nutrients to the parasite.

Thus, broomrape is a hemibiotrophic mixed type of sunflower infection. Initially, it develops as a biotroph and, with the help of effectors, suppresses the course of immune reactions and forms haustoria, which at a certain stage of growth break the membrane bags of cells, lead to cytoplasmic leakage and cause the death of root cells as a necrotroph. At the same time, cell death occurs in the form of necrosis due to damage to root cells by intrusive cells of the parasite’s haustoria tip. The intrusive cells of the haustoria tip move to the vessels of the host root, crossing successive tissues of the sunflower root. The destruction of a part of the sunflower cell wall by broomrape enzymes leads to plasmalemma ruptures as a result of turgor pressure. Through these ruptures, the cell contents spill out into the intercellular space and possibly become a substrate for the parasite to feed on.

There are fundamental differences between hemibiotrophs, biotrophs and necrotrophs in many aspects of their pathogenesis. Infection processes, disease histology, infection-related morphogenesis, the nature of effector proteins, the host defense responses elicited, and their strategies for obtaining nutrition differ significantly. These differences may explain why one defense strategy can effectively limit biotrophs and allow necrotrophs to infect plants. Cell death (HR) limits a biotrophic pathogen by eliminating its nutrients, thereby limiting its growth, but can serve as a substrate for necrotrophic pathogens. Thus, biotrophs actively suppress HR, and necrotrophs promote HR-like cell death (Oliver, Ipcho, 2004).

Because biotrophs have developed complex infection strategies aimed at maintaining host viability and preserving the cytoplasmic membrane of the cell, while necrotrophs have developed tactics to disrupt the integrity of the cell wall and plasma membrane and rapidly kill the cell. Defense reactions that occur before or after HR, including the oxidative burst, can also have contrasting protective functions depending on the pathogen’s lifestyle (Jones, Dangl, 2006).

In contrast to necrotrophs, biotrophic pathogens (powdery mildew, rust, corn smut, tomato leaf mold) secrete a limited number of lytic enzymes, usually do not produce toxins, and evade detection or suppress the immune response by manipulating the host’s defenses. These pathogens establish a close relationship with host cells using specialized structures such as haustoria and begin to slowly deplete plant resources, thereby gradually reducing plant yields (Koeck et al., 2011).

Necrotrophs, on the other hand, are facultative saprophytes that actively destroy host tissue with the help of various toxins and enzymes (CWDEs). Soil-transmitted necrotrophs (fungal species of the genera Rhizoctonia, Fusarium and Colletotrichum and bacterial species of Streptomyces and Ralstonia, as well as the oomycete Pythium) are monocyclic pathogens that produce oligosaccharides in the xylem, which reduces water transport and host viability and causes wilting of seedlings before/after germination and causes wilt and root rot diseases. On the contrary, foliar necrotrophs (belonging to the genera Monilinia, Sclerotinia, Botrytis and Alternaria) are polycyclic, which leads to the formation of a large number of spores that are dispersed over a large area. Generally, resistance in plants to highly specialized soil necrotrophs that have a very limited range of hosts is due to individual genes that provide complete immunity, while resistance to foliar necrotrophs that affect many plant species is quantitative, requiring many genes for full resistance (Laluk, Mengiste, 2010).

Compared to biotrophs and necrotrophs, hemibiotrophic pathogens are characterized by an initial period of biotrophic infection, followed by a necrotrophic phase with host cell death (Perfect and Green, 2001).

The discovery of the hemibiotrophic process of infection of hybrids by the pathogen Orobanche sumana is changing paradigms in the field of parasitic plant research and paving the way for future understanding and development of sunflower broomrape resistance. It is becoming apparent that the salicylic acid (SA) signaling pathway plays an important role in parasitic plant resistance. SA-dependent reactions associated with massive accumulation of ROS and leading to programmed cell death (PCD) by megaautophagy via hypersensitivity response (HR) are an effective defense against biotrophic and hemibiotrophic pathogens. However, the question of what type of megaautophagy occurs in sunflower root cells during broomrape infection needs to be clarified.

This fact opens up a direction in reducing the infection of sunflower with the broomrape parasite through the induction of systemic acquired resistance (SAR) by substances that cause the accumulation of ROS and cause a localized program of cell death through a hypersensitive response (HR). HR involves the generation of reactive oxygen species (ROS), an increase in intracellular Ca 2+ levels, which is often caused by the activation of members of the leucine-rich receptor (NLR) family of intracellular nucleotide-binding domains.

BTH (S-methyl ester of benzo[1,2,3]thiadiazole-7-carbothioic acid - an analog of salicylic acid), riboflavin, amino acid methionine, vitamin B 1 (thiamine), menadione sulfite sodium (MSB), β-aminobutyric acid (BABA), potassium dihydrogen phosphate, potassium phosphonate, ohcom, silicon, herbicides lactophene, trifluralin and ammonium glufosinate, several bacterial and fungal agents (Pseudomonas fluorescens WCS374, Serratia plymuthica ICI270, Bacillus mycoides) are compounds that induce ROS signaling systems and trigger priming of defense responses and induce systemic resistance against pathogens for the treatment of viral, bacterial and phytoplasmic diseases and control of parasitic plants that are difficult to control by traditional chemical methods (Frąckowiak et al., 2019).

Protection of sunflower plants from the broomrape parasite can be activated by chemical or biological means. When infected, necrotrophic and biotrophic pathogens have different effects on the defense mechanisms of the cell, including the manifestation of the autophagy process in the cell. Biotrophs actively inhibit HR cell death, while necrotrophs promote HR-like cell death. Necrotrophic fungi have developed mechanisms to capture and induce plant PCD for their own benefit by colonizing and feeding on dead plant tissues. By using preparations from necrotrophic fungi that cause ROS accumulation and lead to the death of affected cells (PCD) at the site of infection, it is possible to induce systemic acquired resistance (SAR) and activate the immune responses of the culture suppressed by biotrophic pathogens.

Lignin, suberin, and callose are common secondary metabolites that form the secondary cell wall of vascular plants and may strengthen its structure by increasing stiffness at the site of pathogen infection, which confers sunflower resistance to the lupus parasite. Biosynthesis and deposition of lignin or lignin-like phenolic polymers, suberin and callose in cell walls can be rapidly induced chemically and biologically in response to biotic and abiotic stresses and structural damage. In particular, various metabolic enzymes in the lignin biosynthesis pathway are required for resistance against various pathogens. Most of these protective responses occur as early as 1 day after infection and peak on day 3 (Al-Khayri et al., 2023).

Lignification, as a rule, begins during the formation of the secondary cell wall. Usually, plant cells are devoid of lignin during the vegetative stage. As soon as the plant begins to bloom, lignin is produced. Parenchyma cells begin to strengthen their secondary cell walls by producing lignin. Therefore, parenchyma turns into sclerenchyma, a tissue designed for mechanical support. Biotic and/or abiotic stress can also induce lignification in cell walls that normally do not lignify in the absence of stress (Wang et al., 2013).

## Conclusion

It was believed that the broomrape belongs to biotrophic parasites and penetration into the tissues of sunflower roots occurs intracellularly. However, our studies of the behavior of sunflower root cells during the penetration of the pathogen using microscopy methods showed that haustoria broomrape penetrates living tissues of the sunflower root as a result of the degradation of the host cell wall and the formation of a membrane bag. However, at the further stage of infection with the broomrape, when the haustoria penetrates into the middle of the cells and the formation of a membrane bag and the subsequent growth of invasive parasitic structures, membrane ruptures and cytoplasm leakage are observed, which indicates a hemibiotrophic process of sunflower pathogen infection, not a biotrophic one. This discovery is a paradigm shift in the field of parasitic plant research and paves the way for future understanding and development of sunflower resistance to broomrape. It is becoming clear that the salicylic acid (SA) signaling pathway plays an important role in resistance to parasitic plants. This fact opens a direction in reducing the infection of sunflower by the broomrape parasite through the induction of systemic acquired resistance (SAR) by drugs that cause the formation of reactive oxygen species and trigger the protective reactions of plants through the programmed death of cells at the site of infection and cause necrosis of the pathogen.

### Compliance with ethical standards

This article does not contain any research using animals or humans as subjects.

## Notes

https://www.biorxiv.org/content/10.1101/2023.07.24.550254v1

https://bioone.org/journals/the-arabidopsis-book/volume-2010/issue-8/tab.0136/Necrotroph-Attacks-on-Plants-Wanton-Destruction-or-Covert-Extortion/10.1199/tab.0136.full

https://www.frontiersin.org/journals/plant-science/articles/10.3389/fpls.2019.01056/full

### Competing Interest Statement

The authors have declared no competing interest.

